# Locus architecture affects mRNA expression levels in Drosophila embryos

**DOI:** 10.1101/005173

**Authors:** Tara Lydiard-Martin, Meghan Bragdon, Kelly B. Eckenrode, Zeba Wunderlich, Angela H. DePace

## Abstract

Structural variation in the genome is common due to insertions, deletions, duplications and rearrangements. However, little is known about the ways structural variants impact gene expression. Developmental genes are controlled by multiple regulatory sequence elements scattered over thousands of bases; developmental loci are therefore a good model to test the functional impact of structural variation on gene expression. Here, we measured the effect of rearranging two developmental enhancers from the *even-skipped (eve)* locus in *Drosophila melanogaster* blastoderm embryos. We systematically varied orientation, order, and spacing of the enhancers in transgenic reporter constructs and measured expression quantitatively at single cell resolution in whole embryos to detect changes in both level and position of expression. We found that the position of expression was robust to changes in locus organization, but levels of expression were highly sensitive to the spacing between enhancers and order relative to the promoter. Our data demonstrate that changes in locus architecture can dramatically impact levels of gene expression. To quantitatively predict gene expression from sequence, we must therefore consider how information is integrated both within enhancers and across gene loci.

## Introduction

How do changes in regulatory DNA sequence impact gene expression? This question is critical for understanding metazoan development, disease and evolution because precise control of gene expression is necessary for the differentiation and function of metazoan cells. Mis-regulation is increasingly implicated in a broad range of disease states (Karczewski et al. 2013; Maurano et al. 2012), and changes in gene expression underlie some morphological differences between animal species (Wittkopp et al. 2009; Frankel et al. 2011; Mallarino et al. 2011; Manceau et al. 2011; Jones et al. 2012). Natural variation in regulatory DNA is common (Mu et al. 2011; Mackay et al. 2012), but not all changes in regulatory sequence have functional consequences (Romano and Wray 2003; Hare et al. 2008; Swanson et al. 2011). A central challenge is to learn which and to what extent regulatory sequence variants alter gene expression.

Many classes of *cis*-regulatory elements that influence metazoan gene expression have been identified, including enhancers, silencers, insulators and targeting sequences (Maston et al. 2006). Cell type specific expression is primarily directed by enhancers that integrate information from multiple DNA-bound transcription factors (TFs) to produce a specific expression pattern (reviewed in Bulger and Groudine 2010). These short (∼1 kb) sequences can be located upstream, downstream, or within introns of their target gene. Many genes, particularly key developmental TFs, are regulated by several enhancers that together direct the total gene expression pattern (Levine 2010; de Laat and Duboule 2013). Accordingly, mutation or loss of enhancer sequences can have phenotypic consequences (VanderMeer and Ahituv 2011; Dunipace et al. 2011; Kim et al. 2014).

Natural variation in regulatory sequence spans multiple length scales, from single nucleotide polymorphisms (SNPs) to structural variants such as insertions, deletions, duplications, inversions, and translocations that can range in size from 1-10 bp “micro-indels” to 1Mb (|Sudmant et al. 2010; 1000 Genomes Project Consortium et al. 2010; Pang et al. 2010). In humans, structural variation is estimated to account for more than 10 times as much genomic variation between individuals as SNPs (Pang et al. 2010). Specific examples of structural variants have been associated with disease (reviewed in Kleinjan and Coutinho 2009) and morphological evolution (e.g., Jones et al. 2012). Structural variants appear to be under strong purifying selective pressure; structural variants in non-coding sequences are selected against more strongly than non-synonymous base substitutions in coding sequences (Zichner et al. 2013).

Despite the prevalence of structural variation, the consequences of large-scale regulatory rearrangements for gene expression have not been systematically studied. Many studies of regulatory sequence variation have focused on the functional impact of SNPs and small indels, either by directed mutagenesis (Thanos and Maniatis 1995; Arnosti et al. 1996; Swanson et al. 2010), or systematic characterization of enhancer variant libraries (Erceg et al. 2014; Melnikov et al. 2012; Kwasnieski et al. 2012; Smith et al. 2013; White et al. 2013). These studies have elucidated how sequence changes within an enhancer impact its regulatory function. Structural variants, meanwhile, may influence the expression of a gene by changing the relative contributions from different enhancers without altering the individual enhancers themselves. Most simply, deleting enhancers can disrupt gene expression (Ludwig et al. 2005; Guenther et al. 2008; Chan et al. 2010; Dunipace et al. 2011; Montavon et al. 2011; McCarroll et al. 2008). Enhancer duplications also impact gene expression, but in unpredictable ways (Klopocki et al. 2008). Rearrangements that move enhancers relative to one another may also alter expression if their bound TFs interact (Small et al. 1993; Kim et al. 2013). Finally, structural variants might disrupt the 3D structure of a locus, which changes during development (Kagey et al. 2010; Phillips-Cremins et al. 2013) and is important for the regulation of gene expression (Deng et al. 2012; Dekker et al. 2013).

To investigate how structural variants impact gene expression, we created a set of reporter constructs in which we systematically varied the orientation, order and spacing between two enhancers. TFs are known to interact through short-range and long-range repression mechanisms (Gray and Levine 1996; Courey and Jia 2001; Li and Arnosti 2011), we therefore tested a series of distances between enhancers spanning 0 to 1000 bp. We chose to conduct this study in *Drosophila melanogaster* blastoderm embryos because 1) we could use two well-characterized enhancers from the highly studied *even-skipped* (*eve*) locus (Fujioka et al. 1999; Clyde et al. 2003; Struffi et al. 2011); 2) readily integrate our reporters *in vivo* (Groth et al. 2004); and 3) make quantitative measurements of expression at cellular resolution using fluorescent imaging (Luengo Hendriks et al. 2006; Fowlkes et al. 2008; Wunderlich et al. 2014). This powerful system allowed us to quantitatively probe enhancer activity in the full range of cell types present in developing embryos.

Our results demonstrate that structural variants can have a strong effect on gene expression level. First, contrary to the classic definition of enhancers, we found that levels of expression driven by single enhancers vary with orientation and distance to the promoter; the magnitude and direction of this effect was enhancer-specific. Second, in configurations containing two enhancers, expression pattern position was largely maintained but levels of expression varied by nearly 8-fold depending on the orientations, order and spacing of the enhancers relative to one another and the promoter. Third, we found that output driven by two enhancers is not equivalent to additive output from the two component enhancers, even when they are separated by a 1000 bp neutral spacer sequence; this indicates enhancers can interact at a much longer range than previously reported. Taken together, our results suggest that structural variants that alter locus architecture are likely to have a substantial impact on gene expression levels. These results emphasize that in order to quantitatively predict gene expression from sequence we must consider how information is integrated at multiple scales—both within enhancers and across gene loci.

## Results

We chose two well-characterized enhancers from the *eve* locus for our study. *eve* is expressed in 7 stripes along the anterior posterior axis; these stripes are controlled by 5 separate enhancers (Fujioka et al. 1999). We chose eve 3/7 (which drives expression of stripes 3 and 7) and eve 4/6 (which drives expression of stripes 4 and 6). These two enhancers share regulators (Clyde et al. 2003) and are normally located on opposite sides of the locus (Fig. 1A). We engineered various arrangements of these two enhancers to each other and the promoter using typical reporter constructs that contain the *eve* basal promoter driving expression of lacZ (Hare et al. 2008). We integrated these reporters into the same genomic location using the phiC31 site-directed integration system (Groth et al. 2004; Fish et al. 2007). When spacer sequence was required, we used portions of the lacZ coding sequence chosen to minimize predicted binding sites for the regulators of these two enhancers (Supplemental Fig. S1 and S2).

**Figure 1:**
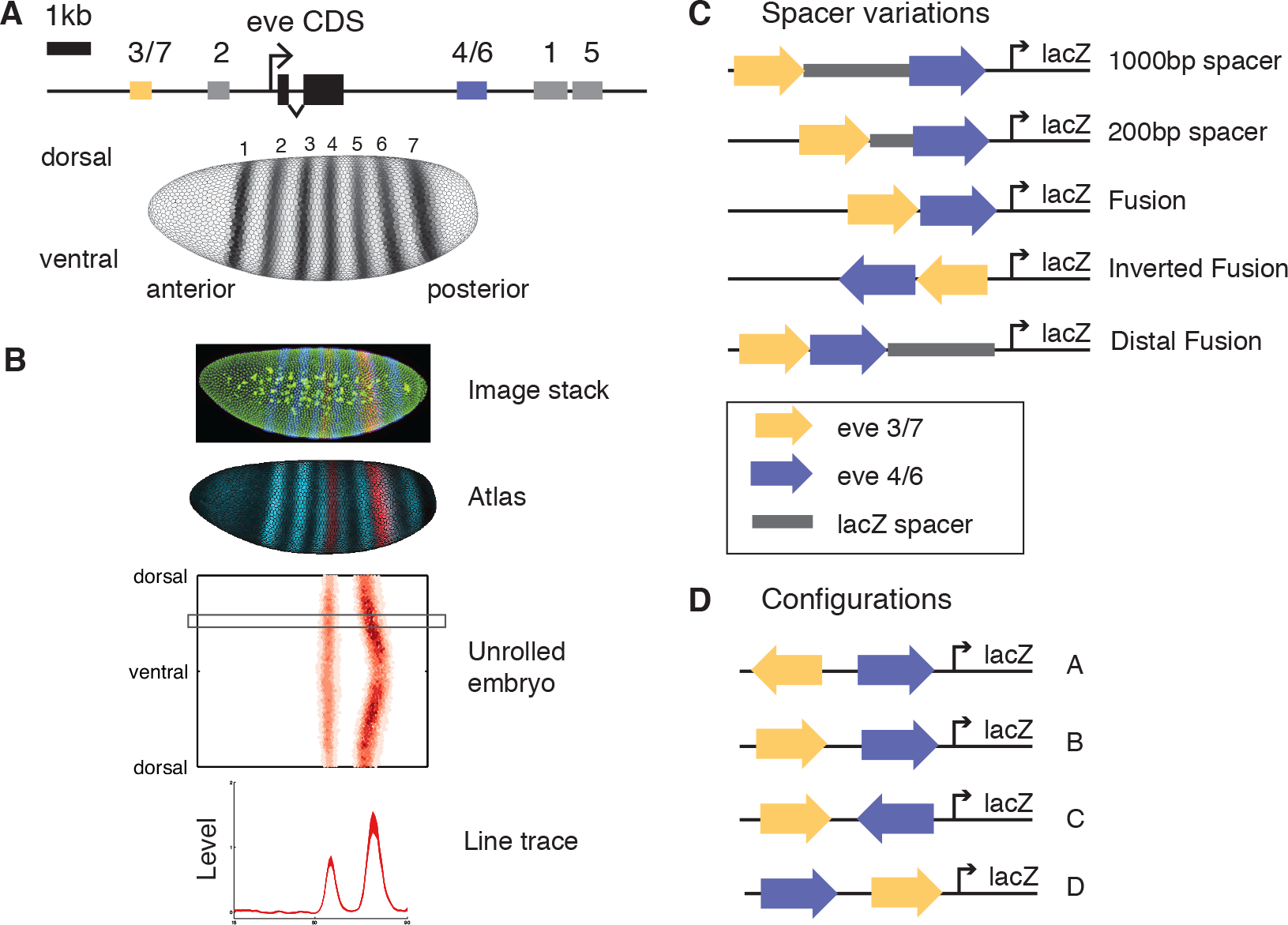
Synthetic reporters containing two enhancers from the *eve* locus test functional consequences of enhancer rearrangements. A) The *eve* locus contains five stripe enhancers encoding the seven stripe pattern of expression in blastoderm embryos. B) We stained embryos for a reporter gene (red) using fluorescent *in situ* hybridization, and collected image stacks through the entire embryo. We computationally segmented embryos and extracted fluorescence values for each cell, then aligned embryos to an average morphological framework to generate an atlas of average expression patterns (see Materials and Methods). During the hour of development under study the cells are in a sheet on the surface of the embryo and can be represented in 2D as an unrolled cylindrical projection. For simplicity, in most figures we show a subset of our data taken from a line trace through the lateral side of the embryo (grey box). C) We tested synthetic arrangements of two enhancers with different length and positioning of spacers. D) We also tested different configurations of the two enhancers that cover all possible junctions between the two.

### Enhancer distance and orientation relative to the promoter affect target gene expression quantitatively

We first measured the effect of changing a single enhancer’s distance and orientation from the promoter. We cloned the minimal eve 3/7 (511 bp; Small et al. 1996) and eve 4/6 (800 bp; Fujioka et al. 1999) enhancers at three positions (0 bp, 500 bp, and 1000 bp upstream of the promoter) and in two orientations (either the endogenous orientation, or reversed). We measured the expression from each reporter construct in blastoderm embryos using fluorescent *in situ* hybridization against the lacZ reporter gene and an endogenously expressed fiduciary marker, *fushi-tarazu* (*ftz*). To normalize levels of expression across reporter constructs we co-stained reporter lines with the endogenous gene *huckebein* (*hkb*) in the same channel as lacZ (Wunderlich et al. 2014). We imaged entire embryos at cellular resolution and assembled our data into a gene expression atlas, which contains average levels of expression for each gene in each cell for six time points during the hour prior to gastrulation (Luengo Hendriks et al. 2006; Fowlkes et al. 2008). For simplicity, in most figures we show a lateral line trace—the moving average of expression level for a five nuclei wide dorsal-ventral (D/V) strip along the anterior-posterior (A/P) axis—for the third time point. The full dataset is publicly available at (depace.med.harvard.edu).

We anticipated that these constructs would merely serve as controls for more complex rearrangements of two enhancers relative to one another, but we found that rearrangements of single enhancers have significant effects on the level, but not the position, of expression. Expression level varies by as much as 2-fold across the constructs we tested, both in terms of overall level of expression and the relative expression of the two stripes (Fig. 2 and Supplemental Fig. S3). For eve 3/7, expression generally decreases as the enhancer moves away from the promoter, but in the reverse orientation this relationship is not monotonic. In contrast, for eve 4/6, expression increases as the enhancer moves away from the promoter. These results demonstrate that there is a complex relationship between expression level and enhancer position and orientation relative to the promoter.

**Figure 2:**
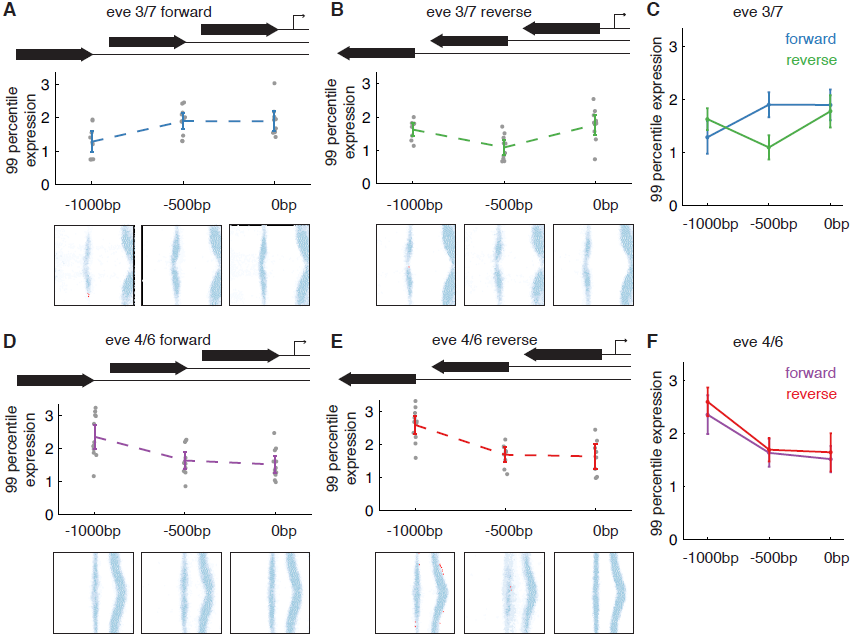
Expression levels driven by the eve 3/7 and 4/6 enhancers depend on enhancer position and orientation relative to the promoter. We measured expression driven by the eve 3/7 (A, B) and eve 4/6 (D, E) enhancers at three distances from the promoter and two orientations as indicated in schematics at top of each panel. We also overlay the measurements for both orientations in order to see the influence of orientation (C, F). We use 99 percentile expression in the trunk (0.2-0.9 egg length) to estimate the level of expression driven by each construct. Expression values were normalized by co-staining with endogenous *hkb* (see Materials and Methods) to enable comparison across transgenic lines. Individual embryos are shown as grey dots; black bars indicate the mean and 95% confidence interval of the mean. We observe significant differences in expression dependent on distance and orientation. We also thresholded gene expression in the embryos to test whether the position of expression changed. We show an unrolled embryo view for each distance with the percentage of embryos in which a cell expresses the reporter plotted in blue. Cells that were significantly different from the reference line (0 bp from promoter in forward orientation) are plotted in red (p < 0.05, Fisher’s Exact Test with permutation to control for multiple hypothesis testing). Position does not change for most lines. The most extreme position shift is a narrowing of the stripes in reverse orientation eve 4/6 at 1000 bp from promoter (E).

Despite the changes in expression level, the set of cells expressing the reporter gene was largely consistent in different transgenic lines. After thresholding the gene expression patterns (see Methods), we identified only a handful of cells with statistically significant changes in expression (Fig. 2). For eve 3/7, these cells are associated with variation in expression along the D/V axis. For eve 4/6, the enhancer drives slightly narrower stripes in the reverse orientation at −1000 bp than when it is in the forward orientation adjacent to the promoter. The qualitative similarity of the expression patterns is consistent with previous studies which found that the stripe enhancers drove expression in the appropriate cells even when moved from their endogenous context to a reporter (Small et al. 1992; 1996; Fujioka et al. 1999). These studies used p-element insertions and were therefore limited to qualitative techniques that could accurately measure expression position, but not level. To fully capture the effects of locus organization on gene expression, cellular resolution quantitative methods are required.

### Enhancers do not act independently even when separated by a large spacer sequence

We next tested how arrangement and spacing of two enhancers relative to one another and the promoter influences expression pattern. We created a set of constructs using eve 3/7, eve 4/6, and spacer sequences to systematically test the influence of spacing between enhancers (Fig. 1C). For each spacing we tested several arrangements, labeled A-D, with the spacing indicated by a subscript (Fig. 1D). Our choice of spacing was based on the distance over which short-range repressors can act, because each *eve* enhancer employs short-range repressors to direct stripe expression (Clyde et al. 2003; Struffi et al. 2011). Short-range repressors bound at one enhancer are capable of disrupting the activity of another enhancer only if placed within 150 bp (Fakhouri et al. 2010). We therefore created constructs where the two enhancers are separated by 1000 bp, 200 bp, and 0 bp.

The eve 3/7 and eve 4/6 enhancers are normally on opposite sides of the gene, separated by approximately 9 kb, and thought to act additively (Maeda and Karch 2011). We hypothesized that they would still act additively when both are placed upstream of a reporter gene if separated by a sufficiently large neutral spacer sequence. To test this hypothesis, we created a set of four constructs containing the two enhancers upstream of the promoter with a 1000 bp spacer between them, where the orientation and order of the enhancers varies relative to one another and the promoter. Our null expectation was that the output of the two enhancers would simply add together; we calculated this null expectation by adding the expression patterns we measured for each single component enhancer at the properly controlled position and orientation. Comparing the expression patterns driven by our constructs to the null expectation clearly revealed non-additive behavior that depended on the orientation and arrangement of the enhancers (Fig. 3). The largest discrepancy was for D_1000_, where expression of stripes 3 and 7 was virtually abolished. In A_1000_, B_1000,_ and C_1000_ stripe 3 expression increased while stripe 7 did not change, indicating that the two stripes do not always change expression in a coordinated way. The eve 4/6 enhancer had lower than expected expression in A_1000_, and B_1000,_ but increased slightly in C_1000_. We conclude that enhancer function is sensitive to the presence of other enhancers in the locus and that the underlying mechanism is affected by the position and orientation of the enhancers relative to one another.

**Figure 3:**
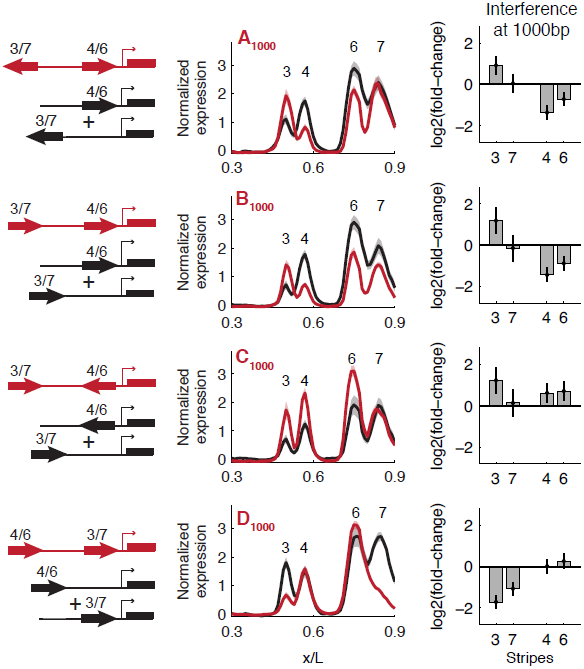
Some configurations of enhancers separated by 1000 bp produce non-additive expression. We created a set of four constructs containing the two enhancers in different orientations and orders relative to one another with a 1000 bp spacer sequence between them. We compared each construct to a null hypothesis of additive activity, as illustrated in the schematics on the left. Normalized expression as a function of fraction of egg length (x/L) is shown for lateral line traces of test constructs (red), and the null hypothesis (black). Shadows indicate standard error of the mean (SEM). We also measured the fold-change in mean expression of each stripe relative to the single enhancer controls. We plot the log_2_(fold-change) so that increases and decreases in expression appear with comparable magnitudes. Error bars indicate 95% confidence interval of the mean.

To define the range of influence of enhancers on one another’s activity, we moved the enhancers closer to one another in the same four configurations. With a 200 bp spacer, we observed additional changes in the level of target gene expression compared to constructs with a 1000 bp spacer (Fig. 4). Specifically, the reduced expression of eve 4/6 was even more pronounced in A_200_, and B_200_, while D_200_ showed no additional interaction between the two enhancers. In all four configurations, the enhancer closest to the promoter drove lower levels of expression. We conclude that enhancers influence each other’s output when they are separated by distances of 200-1000 bp, a much longer distance than previously described for interactions between eve 3/7 and eve 2 (Small et al. 1993).

**Figure 4:**
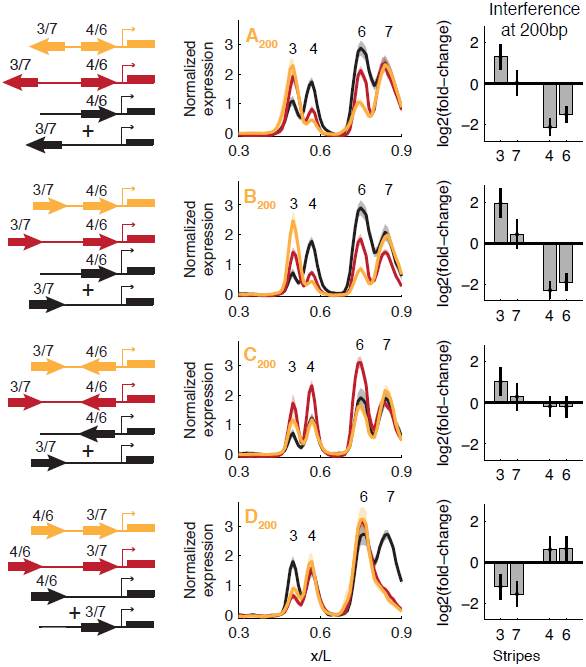
Enhancers influence each other’s level when in close proximity. We compared expression from configurations containing a 200 bp spacer to 1000 bp spacers and the additive null hypothesis. Normalized expression as a function of fraction of egg length (x/L) is shown for lateral line traces for configurations with a 200 bp spacer (yellow), 1000 bp spacer (red), and the null hypothesis (black). Shadows indicate SEM. We also plot the log_2_(fold-change) in mean expression of each stripe relative to the single enhancer control for the 200 bp spacer constructs. Expression of stripes 4 and 6 are consistently reduced in configurations A, B and C relative to both the 1000 bp spacer version and single enhancer controls.

### Fused enhancers direct expression patterns only slightly shifted in position

To quantify the influence of short-range interactions between the two enhancers, we fused them together. We expected interactions between the component enhancers to occur at the junctions, due to local interactions between TFs such as short-range repression and cooperative binding; the four configurations represent all possible junctions between the two enhancers. Previous studies have indicated that short range repression is able to quench activation for up to 150 bp on either side of the repressor binding site (Fakhouri et al. 2010; Gray and Levine 1996). Cooperative binding between TFs operates over an even shorter length scale (Crocker et al. 2008; Hanes et al. 1994). Because eve 3/7 and eve 4/6 stripe boundaries are regulated by the same pair of short-range repressors (Clyde et al. 2003; Struffi et al. 2011; Supplemental Fig. S4), we expected that these TFs would act across the junctions of the fused enhancers, thus changing the position of the stripe boundaries driven by this set of reporters (map of TF binding sites in Supplemental Fig. S1 and S2).

However, when the eve 3/7 and eve 4/6 enhancers were fused together the position of the stripe boundaries changed only slightly in two configurations; expression level was affected in three configurations (Fig. 5). We compared the expression driven by enhancers separated by 200 bp to those directly juxtaposed in order to compare the influence of locus arrangement to the influence of short-range transcription factor interactions. A_fusion_ exhibited no additional changes in expression level or shifts in expression pattern boundaries. The only expression domain that moved was stripe 7; it shifted anteriorly in C_fusion_ (∼1 nucleus width, Supplemental Fig. S5). Expression levels of stripes 4 and 6 were slightly higher in B_fusion_ and C_fusion_, and expression in the region of stripe 7 was substantially higher in B_fusion_ and slightly increased in D_fusion_. We conclude that interactions between these two enhancers, even at short distances, predominantly affect expression level, rather than the boundaries of expression patterns.

**Figure 5:**
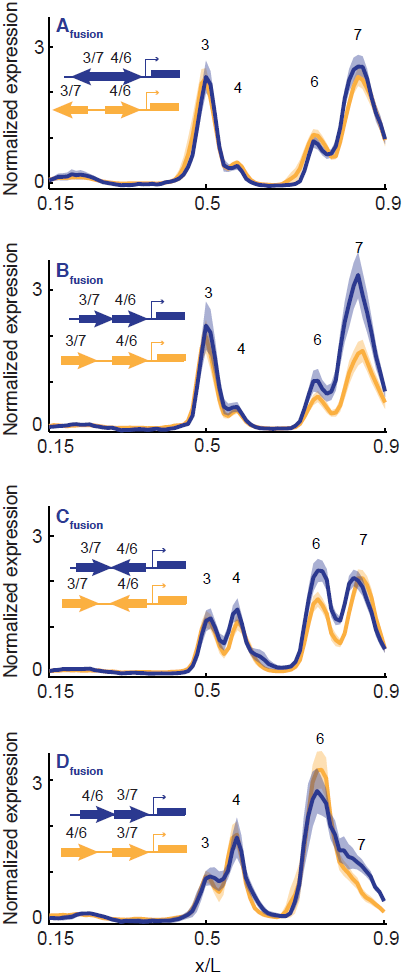
Local transcription factor interactions have a minor effect on expression from fused enhancers. We compared fused enhancers (blue) to the same configuration with a 200 bp spacer (yellow) to estimate the influence of local interactions between transcription factors bound at the junction on expression. B_fusion_ shows an increase in stripe 7 accompanied by an anterior shift in expression. C_fusion_ shows the same shift in stripe 7 without increased expression. Shadows indicate SEM.

### Levels of gene expression depend on order of enhancers relative to the promoter

Our data suggest that order of the enhancers relative to the promoter has a significant influence on expression level. The most consistent effect of the enhancer arrangement on target gene expression was a reduction in the level of expression driven by the promoter proximal enhancer. We tested the hypothesis that order relative to the promoter influences the level of expression driven by each enhancer by inverting entire fusions.

Inverting the fusions switched the relative levels of expression driven by the two enhancers. We also observed changes in the relative expression of two stripes driven by the same enhancer (Fig. 6). B_inverted_ retained high levels of stripe 3 expression, even as stripe 7 expression was reduced nearly 4-fold. We conclude that order of enhancers relative to the promoter has a strong effect on levels of expression, but that other characteristics, such as orientation, also influence level. In combination with our findings from the single enhancer experiments, these results suggest that distance from the promoter needs to be considered both within enhancers where it manifests as orientation dependence, and across the locus.

**Figure 6:**
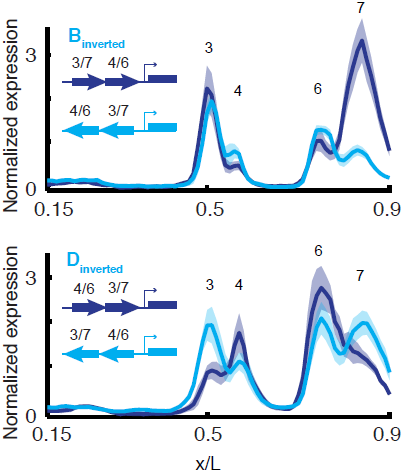
Relative proximity to the promoter influences level of expression driven by enhancers. We compared the expression of two fusions (dark blue) to a complete inversion of the entire fusion construct (light blue). In both cases, we see a reversal of the relative levels driven by each component enhancer. Stripes 3 and 7 show large changes in level of expression, while the effect on stripes 4 and 6 is smaller. Shadows indicate SEM.

### Fused enhancers still interact when moved away from the promoter

In all of our constructs one enhancer was immediately adjacent to the promoter. The promoter may exert an influence on enhancer function, either through chromatin or by the basal transcriptional machinery or associated factors. Many promoters have a well positioned nucleosome upstream of the transcription start site (Mavrich et al. 2008), which might occlude portions of the enhancer. Alternatively, the TFs bound at the promoter proximal enhancer could interact directly with the promoter by a different mechanism than when they are farther away. We therefore tested whether moving fused enhancers away from the promoter relieved the repression of the promoter-proximal enhancer.

We found that fusions placed 1000 bp upstream of the promoter still drove the same unequal levels of expression as fusions immediately adjacent to the promoter (Fig. 7). The predominant consequence of moving the fusions away from the promoter was a reduction in the expression in stripes 3 and 7, which is consistent with the observation that the stripe 3/7 enhancer alone had reduced expression when moved away from the promoter. We conclude that depressed expression from the promoter-proximal enhancer does not require a direct juxtaposition of the enhancer and the promoter.

**Figure 7:**
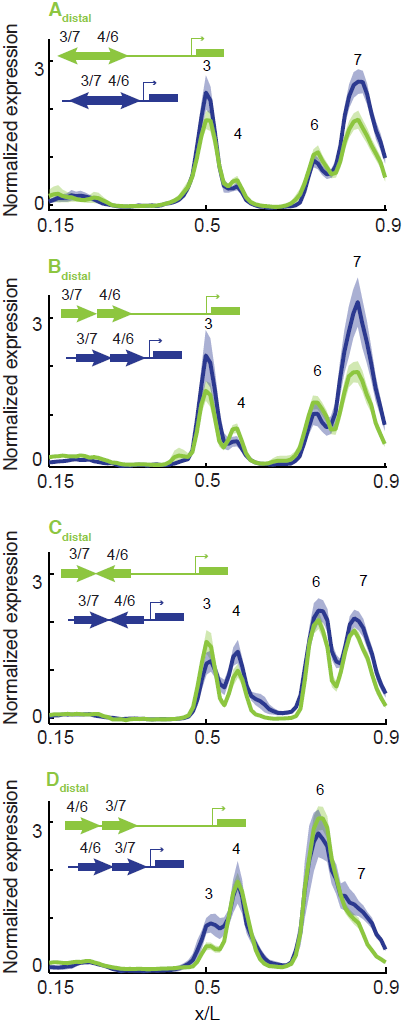
Fused enhancers still interact when moved away from the promoter. We tested the role of the local promoter environment on determining the level of expression of the proximal enhancer by introducing a 1000 bp spacer between fused enhancers and the promoter. Expression of fusions at a distance from the promoter (green) is similar to expression adjacent to the promoter (blue), with the exception of eve 3/7, which is lower in configurations A, B and D, consistent with eve 3/7 having lower expression as it moves away from the promoter. Shadows indicate SEM.

## Discussion

To assess the effect of structural variation on gene regulation, we measured reporter gene expression driven by different arrangements of two *eve* enhancers in *Drosophila melanogaster* blastoderm embryos. We systematically varied orientation, order, and spacing of enhancers and measured expression quantitatively at single cell resolution to detect changes in level and position of expression. This approach allowed us to quantify the influence of locus organization while minimizing changes in sequence content. We found that multiple features of locus organization affect expression level significantly; we found only minor changes to the position of the expression pattern. This partially contradicts the classic definition of enhancers as modular units capable of driving the same expression pattern regardless of orientation or distance from the promoter and independent of the activity of other enhancers in the locus (reviewed in Maston et al. 2006). First, we found levels of expression driven by single enhancers vary with the enhancer’s orientation and distance from the promoter; the direction and strength of this effect is enhancer-specific. Second, we found that in constructs containing two enhancers, levels of expression depend on the order and spacing of the enhancers relative to each other and the promoter. Third, we show that the total expression driven by pairs of enhancers can be non-additive, even when the enhancers are separated by 1000 bp, a much longer range expected for short-range repression mechanisms.

### Distance between enhancer and promoter influences expression level

Here we show that the expression driven by two single *eve* enhancers is sensitive to the enhancer’s position relative to the promoter and this effect is enhancer-specific (Fig. 2B). These results are consistent with experiments using the SV40 enhancer (Wasylyk et al. 1984) and IFN-beta enhanceosome (Nolis et al. 2009), which found that levels of expression driven by these enhancers also depended on distance from the promoter.

One caveat to this experiment is that the sequence adjacent to the 3’ end of the enhancer changes at each distance due to the different lengths of spacer sequence used. It is thus possible that short-range interactions between transcription factors (TFs) bound near the junction between enhancer and spacer sequence are responsible for the distance dependence that we observe. The degeneracy of eukaryotic transcription factor binding motifs (Wunderlich and Mirny 2009) makes it difficult to completely eliminate all TF binding sites in spacer sequences, and the possibility of introducing inappropriate interactions always exists. We examined the predicted TF binding sites in the spacer sequences and found no clear candidates that would explain the observed trends (Supplemental Fig. S1 and S2). The spacer sequences contain few predicted TF binding sites, and there are few activator binding sites near the edges of the enhancers, which we would expect to be most strongly affected by short-range interactions. The introduction of long-range repressor binding sites within the spacer could globally decrease expression (Courey and Jia 2001). However, in this case we would expect the effect of distance to be enhancer-independent since the constructs used the same spacers. Instead, the trends are in opposite directions, which suggests that the distance-dependence is not a function of the spacer sequences.

Enhancer-promoter interactions may differ depending on whether an enhancer is promoter-proximal or acting at a distance. Most promoters include a well positioned nucleosome approximately 180 bp upstream of the transcription start site (TSS) (Mavrich et al. 2008); when enhancers are in close proximity to the promoter this nucleosome may occlude some binding sites. In yeast, where most regulatory sequences are promoter proximal, nucleosome position has a large effect on which TF binding sites are used (Kim and O’Shea 2008; Raveh-Sadka et al. 2012). In addition, the pre-initiation complex (PIC) containing RNA Pol II, general TFs and co-factors forms a large complex spanning ∼100 bp across the TSS, and several components have been found to induce DNA bending (reviewed in Levine et al. 2014). Hence, TFs bound to promoter proximal enhancers can come into direct contact with elements of the PIC (Park and Hong 2012). Conversely, metazoan enhancers commonly act a distance via looping mediated by mediator, cohesin, and TF binding sites in both the enhancer and promoter (Phillips-Cremins et al. 2013; Kagey et al. 2010; Su et al. 1991). For example, expression of the β-globin gene and looping between the locus control region (LCR) and β-globin promoter are eliminated in GATA1 null cells, but tethering of the two elements with an artificial zinc finger enabled looping and rescued transcription (Deng et al. 2012). The fly *sparkling* enhancer contains a “remote control element” which is required for the enhancer to drive activity at a distance of 846 bp, but not when adjacent to the promoter (Swanson et al. 2010). Taken together, these studies support the idea of direct activation when enhancer and promoter are proximal, and a switch to action at a distance mediated by looping.

How might looping result in different levels of expression than direct interaction between enhancers and promoters? It is possible that once looping is established, the interaction between enhancer, bound TFs, and the PIC is the same as when the enhancer is promoter proximal. Thus, the changes in expression level we observe with enhancer-promoter distance may be due to different frequencies or stabilities of enhancer-promoter interactions. However, it is also possible that acting at a distance allows greater conformational freedom and consequently changes the physical interaction of enhancer bound TFs and the promoter in an enhancer-specific manner.

### Orientation of enhancer relative to promoter influences expression level

The level of expression driven by individual enhancers in our study is sensitive to enhancer orientation. This is particularly evident when the eve 3/7 enhancer is −500 bp from the promoter (Fig. 2); at this distance the eve 3/7 enhancer drives significantly different levels of expression in each orientation. The relative levels of each stripe driven by a single enhancer (e.g. stripe 3 and stripe 7) also vary with both distance and orientation (Supplemental Fig. S3). These data demonstrate that the regulatory sequences that generate each stripe are somewhat separable; the stripes need not change in concert. The location of TF binding sites in the enhancer is asymmetric. The orientation dependence may therefore be due to either different TFs coming into contact with TFs bound to the spacer, or the underlying distance-dependence of the TFs interacting with the promoter. Distance-dependent activity for individual binding sites has been demonstrated in both bacteria (Garcia et al. 2012) and yeast (Sharon et al. 2012). Even in these relatively simple systems with single binding sites the distance dependence function is complex. Enhancers contain many TF binding sites, and the aggregate output if each of those binding sites has distance-dependent activity is hard to predict.

In summary, we suggest that the orientation effect is likely due to a combination of asymmetric distribution of binding sites combined with a dependence on distance from the promoter. At minimum, these experiments demonstrate that the information processing in enhancers is asymmetric and highly sensitive to locus context.

### Levels of expression depend on the distance and orientation of enhancers relative to each other and the promoter

We found that the largest impact on level of expression was due to interactions between two enhancers in the same reporter construct. The classic definition of enhancers as autonomous units led us to formulate the null hypothesis that the two enhancers would have additive outputs. Contrary to our expectation, we observed a large non-additive interaction effect on level of expression. Our experiments do not address whether the interaction effect is due to direct physical interaction or indirect interaction, for example, through competition for the promoter. However, we can make some observations about the character of the interaction. The largest effects are correlated with the order of the enhancers relative to the promoter. In general the enhancer closest to the promoter directs lower expression than expected, while the more distant enhancer directs normal or elevated expression (see Figs. 3, 4 and especially 6). The strength of the interaction is dependent on the distance between the enhancers, for 3 of 4 cases. The exception to this rule is configuration D_1000_, in which the repression of eve 3/7 is extremely strong at all distances tested. The magnitude of this effect is much stronger than the effect of short-range interactions between TFs bound at the junctions between fused enhancers (Fig. 5). Finally, we confirmed that the interaction effect is not due solely to one enhancer being directly adjacent to the promoter (Fig. 7).

One possible explanation for the observed interaction effect is the formation of direct physical interactions between the enhancers. Many TFs recruit co-factors and adapters for the explicit purpose of establishing long range interactions with the promoter (Phillips-Cremins et al. 2013; Kagey et al. 2010; Su et al. 1991), and these may target other enhancers as well. The two enhancers we used share the same regulating TFs but produce different positions of expression due to different sensitivities to the repressors *hunchback* (*hb*) and *knirps* (*kni*) (Clyde et al. 2003; Struffi et al. 2011). The maintenance of the stripe positions implies that the two enhancers retain separate information integration functions. This constraint argues against the direct interaction of the two enhancers through the formation of a single large complex.

In addition to activating transcription from the promoter, it has recently been shown that enhancers are themselves transcribed (Kim et al. 2010). Enhancer RNAs (eRNA) are generally short-lived, but a variety of putative functional roles have been assigned to them, including recruitment of co-factors and facilitating looping (reviewed in Lam et al. 2014). In yeast, when two promoters drive expression of a single gene, the upstream promoter is used preferentially because transcription through the downstream promoter disrupts its activity (Hirschman et al. 1988; Iyer and Struhl 1995; Martens et al. 2004). It is possible that the interaction between enhancers that we observe is due to a similar effect in which the eRNA produced by one enhancer interferes with the activity of the other.

An alternate, indirect, form of interaction between enhancers is through chromatin spreading, which is primarily associated with silencing through long-range repression (Courey and Jia 2001; Li and Arnosti 2011). Some enhancers recruit chromatin-modifying enzymes, which alter the chromatin composition of the locus and might produce either silencing or enhancement of nearby enhancers (discussed in Bulger and Groudine 2011). However, this mechanism would be expected to depend only on presence or absence of a second enhancer, not on the relative arrangement of the two. Even if the chromatin spreading was directional, we would expect to see an effect that was more strongly dependent on the orientation of the enhancers rather than order relative to the promoter.

An intriguing possibility is that the order of enhancers relative to the promoter may influence the 3D structure of the locus and thus the efficiency of enhancer-promoter interactions. Numerous studies have found correlations between enhancer-promoter looping and gene expression (Deng et al. 2012; Chopra et al. 2012). In addition, a study of the *hox* locus found that enhancers in the locus formed a set of looped contacts even in a transcriptionally silent state supporting the idea that transcriptionally silent enhancer elements regulate the 3D structure of the locus (Montavon et al. 2011). In our constructs the promoter proximal enhancer may be looped out by the distal enhancer, reducing its expression. However, this explanation does not account for increased expression of the distal enhancer. Most likely we are seeing the combined effects of multiple processes, including regulated looping.

It is important to note that the two enhancers in our study drive expression in different sets of cells. The existence of an interaction effect therefore indicates that even when enhancers are transcriptionally silent they can influence one another’s output. In differentiating cells, enhancers recruit chromatin modifying activity and may interact with basal transcriptional machinery prior to becoming transcriptionally active (Rada-Iglesias et al. 2011; Creyghton et al. 2010). Our data suggest that “poised” enhancers may influence the activity of neighboring regulatory sequences as well.

### Implications for interpreting regulatory sequence variants

Current computational models focus on predicting the activity of single enhancers and do not take locus-level features into account. Single enhancer models are reasonably successful at predicting expression patterns, but do not scale up to the whole locus well (Kim et al. 2013; Samee and Sinha 2014). Using quantitative methods, we have shown that rearrangements of enhancers may affect target gene expression levels, even when binding site content within the enhancer is maintained. In addition, duplications and deletions are likely to have non-additive effects. Our results suggest that including locus-level parameters beyond TF binding will be necessary for accurate predictions.

### Implications for regulatory sequence evolution

Given our results that locus organization can affect expression level, selection for expression level may explain conservation of locus architecture. A recent population genetics study in *Drosophila* found that structural variants in both coding and non-coding sequences showed evidence of strong purifying selection (Zichner et al. 2013). Studies in both vertebrates and insects have identified regions of “micro-synteny” in which recombination events are much lower than expected (Sun et al. 2006; Engström et al. 2007; Cande et al. 2009). These regions are enriched for developmental genes and highly conserved elements, a proxy for enhancers. Together, these observations point to an important role for locus architecture in the function of developmental genes.

## Materials and Methods

### Construction of reporters and transgenic lines

We used RedFly to identify coordinates of the eve stripe 3/7 and stripe 4/6 enhancers (Gallo et al. 2011). The eve_stripe_3 + 7 element is 510 bp (Release 5 coordinates 2R:5863006-5863516) (Small et al. 1996), while the eve_stripe4_6 element is 800 bp (Release 5 coordinates 2R: 5871404-5872203) (Fujioka et al. 1999). Note that the stripe 4/6 enhancer coordinates from REDfly contain an extra 208 bp on the 3’ end compared to the construct tested in Fujioka *et al*. (1999). Enhancers were PCR amplified from genomic DNA from *w*^118^ *Drosophila melanogaster* flies and sequence verified. Enhancers were inserted into the multiple cloning site of the pBOY vector (Hare et al. 2008) using isothermal assembly (Gibson et al. 2009), which leaves scar-less junctions. LacZ spacer sequences were amplified from the pBOY vector. pBOY contains an *eve* core promoter 20 bp downstream of the multiple cloning site that drives an eve/lacZ fusion transcript. The vector also contains an attB site for phiC31 site specific integration (Fish et al. 2007) and the *mini-white* gene for selection of transformants. Each plasmid was injected into attP2 flies (Markstein et al. 2008) by Genetic Services, Inc and transgenic flies were homozygosed using the mini-white eye color marker.

### Embryo collection and *in situ* hybridization

Embryo collection and whole mount *in situ* hybridization was performed as previously described (Luengo Hendriks et al. 2006). Briefly, 0-4hr embryos (25C) were collected, dechorionated in 50% bleach, fixed in a 1:4 mixture of 10% formaldehyde to heptane, and devitellinized in heptane and methanol by shaking. Embryos were post-fixed in formaldehyde and a formaldehyde based hybridization buffer. Hybridizations were performed at 56C with two or three full length cDNA probes: a DIG-labeled probe for *fushi tarazu* (*ftz*), a DNP-labeled lacZ probe and optionally a DNP-labeled probe against *huckebein* (*hkb*). The probes were detected by successive antibody staining using anti-DIG-HRP (anti-DIG-POD; Roche, Basil, Switzerland) and anti-DNP-HRP (Perkin-Elmer TSA-kit, Waltham, MA, USA), and labeled by reactions with coumarin-and Cy3-tyramide (Perkin-Elmer). Embryos were treated with RNase and incubated with Sytox Green (Invitrogen, Carlsbad, CA, USA) to stain nuclei. Finally, embryos were dehydrated in ethanol and mounted in DePex (Electron Microscopy Sciences, Hatfield, PA, USA), using #1 coverslips to form a bridge to preserve 3D embryo morphology.

### Imaging and image processing

Embryos were imaged and computationally segmented for further analysis (Luengo Hendriks et al. 2006; Fowlkes et al. 2008). A three-dimensional image stack of each embryo was acquired on a Zeiss LSM Z10 with a plan-apochromat 20 × 0.8 NA objective using 2-photon microscopy. Embryos were binned into six time points of approximately 10 minute windows using the extent of membrane invagination under phase-microscopy as a morphological marker. Time points correspond to 0-3%, 4-8%, 9-25%, 26-50%, 51-75% and 76-100% membrane invagination along the side of the embryo that has progressed most. Image files were processed into PointCloud representations containing the coordinates and fluorescence levels for each nucleus. Using the *ftz* fiduciary marker, PointClouds were registered to an average morphological template to create a gene expression atlas, a summary text file containing the normalized expression level for each reporter construct in each nucleus at each time point.

### *hkb* normalization

Normalization to a *hkb* co-stain was performed to test the variation in absolute levels of expression across reporters (Wunderlich et al. 2014). Embryos were stained with a mixture of *lacZ*-DNP and *hkb*-DNP probe. Stains were done in two batches: the first batch contained all single enhancer control lines; the second batch contained all two enhancer constructs and two single enhancer control lines to allow comparison between batches. For each embryo, background was calculated as the mode of the fluorescence distribution. After subtracting background, mean *hkb* fluorescence was calculated as the geometric mean of the anterior and posterior expression domains. We noted that eve stripe 7 overlaps slightly with the posterior expression domain of *hkb*, and so chose to use the geometric mean of anterior and posterior rather than solely the posterior domain as in (Wunderlich et al. 2014) to limit the impact of overlapping expression. The fluorescence in each nucleus was then divided by the mean *hkb* fluorescence to yield a normalized expression level.

### Data analysis and visualization

Extraction of lateral line traces, and embryo thresholding were performed in MATLAB using the PointCloud Toolbox (http://bdtnp.lbl.gov/Fly-Net/bioimaging.jsp?w=analysis) and custom scripts. Briefly, lateral line traces are a smoothed moving window average over a 1/16th DV strip (about 5 nuclei wide) along the left side of the embryo. Lateral line traces were taken for each individual embryo after atlas registration and the mean and standard error of the mean was calculated for each point along the AP-axis. To measure positional variation in Fig. 2 individual embryos were thresholded into on/off using the mode + standard deviation of expression values in the trunk (0.2-0.9 egg length) as the threshold. For each cell, the distribution of on/off calls was compared using fexact.m which computes Fisher’s Exact Test with permutation (10 times) to control for multiple hypothesis testing.

To find predicted TF binding sites shown in Supplemental Figures S1 and S2, we used Patser (http://stormo.wustl.edu/software.html), with PWMs as listed in Supplemental Table S1. Background GC content was set to 0.406, a P-value limit of 0.001 was used. We plotted the predicted binding sites using InSite, an interactive tool developed by Miriah Meyer (http://www.cs.utah.edu/∼miriah/insite/).

## Acknowledgments

The authors wish to thank Ben Vincent, Clarissa Scholes, and Max Staller for insightful discussions and careful editing of the manuscript. We also wish to thank Saurabh Sinha, and Md Abul Hassan Samee for sharing modeling predictions; these results shaped our thinking throughout the project.

